# Risk variant in miR-544a binding site in *CCDC170* is associated with osteoporosis

**DOI:** 10.1101/595918

**Authors:** Xinhong Liu, Yang Wang, Qin Tan, Yunqi He, Xin Zhao

## Abstract

MicroRNAs (miRNAs) play essential roles in regulating bone formation and homeostasis. Genomic variations within miRNA target sites may therefore be important sources of genetic differences in osteoporosis risk. To investigate this possibility, we searched for miRNA recognition sites within genes using the TargetScan, miRNASNP and miRbase databases. In this study, we showed that miR-544a differentially regulated the allele variants of rs6932603 in the *CCDC170* 3 ′ untranslated (3′-UTR). We also showed that miR-544a suppressed osteogenesis and promoted osteoclastogenesis by regulating the expressions of *Runx2*, *Osterix, Alp, Col1a1*, *Osteopontin* (*OPN*) *,Osteocalcin* (*OCN*) and *Osteoprotegerin* (*OPG*). Our results suggest that allele-specific regulation of *CCDC170* by miR-544a explains the observed disease risk, and provides a potential therapeutic target for osteoporosis therapy.

## Introduction

Osteoporosis is a common disease characterized by low bone mass and defects in the microarchitecture of bone tissue, which impairs bone strength and leads to an increased risk of fragility fracture responses[1]. With increased age and changes in body hormones, the body’s bone mass decreases. When bone mass decreases and bone fragility increases, fractures and osteoporosis will eventually occur[2]. Discovering genetic variation loci and clarifying their biological functions in bone mineral density (BMD) variations are important to understanding the etiology of osteoporosis and developing new approaches to screen, prevent, and treat osteoporosi [3].

MicroRNAs (miRNAs) are single-stranded noncoding RNA molecules of approximately 21–23 nucleotides long, which participate in posttranscriptional regulation of gene expression by directly binding to the 3′ untranslated region (3′-UTR) of target mRNAs[4–7]. MiRNAs play important roles in the occurrence of various diseases, including osteoporosis[8]. In recent years, multiple studies have shown that miRNAs can regulate bone formation and remodeling[9–11].

Single-nucleotide polymorphism (SNP)-based genome-wide association studies (GWASs) have already identified approximately 100 BMD-associated loci[12], including *CCDC170*. Multiple GWASs have shown that *CCDC170* is one of the genes most strongly linked to BMD across populations[13–15]. Recent study have found that multiple SNPs in *CCDC170* are associated with BMD, including rs6929137, rs371804, rs3734805, rs6932260, rs6904261, rs6932603, rs9383935, rs9383589, rs3734806 and rs3757322[16]. The functions of most of these SNPs in vivo and in vitro remain unverified.

In the present study, we examined the role of *CCDC170* and miR-544a in osteosarcoma. We found that miR-544a differentially bound to the GWAS lead SNP rs6932603 (C-T) (in the *CCTC170* 3′-UTR) and inhibited osteogenesis by regulating the expressions of *RUNX2*, *OSX*, *ALP*, *COL1A1*, *Osteopontin* (*OPN*), *Osteocalcin* (*OCN*) and *Osteoprotegerin* (*OPG*). In summary, our study showed that rs6932603 may act as an etiological factor of osteoporosis and could be a possible target in treating osteoporosis.

## Materials and methods

### Cell cultures

The human embryonic kidney cell line, HEK293T, was obtained from the Cell Bank of Wuhan University (Wuhan, China). The human osteosarcoma cell line, U2OS, was purchased from the Cell Bank of the Chinese Academy of Sciences (Shanghai, China). HEK293T cells were cultured in Dulbecco’s modified Eagle’s medium (DMEM) (HyClone, Waltham, MA, USA), and U-2OS cells were grown in McCoy’s 5A medium (HyClone). Media were supplemented with 10% fetal bovine serum (Invitrogen, Carlsbad, CA, USA), 100 U/ml penicillin, and 100 μg/ml streptomycin (Invitrogen). All cells were maintained at 37°C in a humidified incubator at 5% CO2.

### Cell transfection and luciferase reporter assay

The rs6932603 C and T alleles were amplified and cloned into psi-check-2 vectors by inserting them between the SgfI/NotI sites. Table 1 shows the sequences cloned into the vectors. Cells were si-transfected with the rs6932603 C/T allele psi-check-2 vector or cotransfected with rs6932603C/T allele psi-check-2, miR-544a mimics, NC mimics, miR-544a inhibitors or NC inhibitors at 20 nM using Lipofectamine 3000 (Invitrogen) with serum-free media for 48h per the manufacturer’s instructions. The luciferase reporter assay was performed using the Dual-Luciferase Reporter Assay (Yeasen); the relative luciferase activity was calculated after transfection for 48h by normalizing the Firefly luminescence to that of Renilla. Experiments were performed in triplicate, and all experiments were repeated three times. Table 1 lists the primer sequences.

**Table 1.**
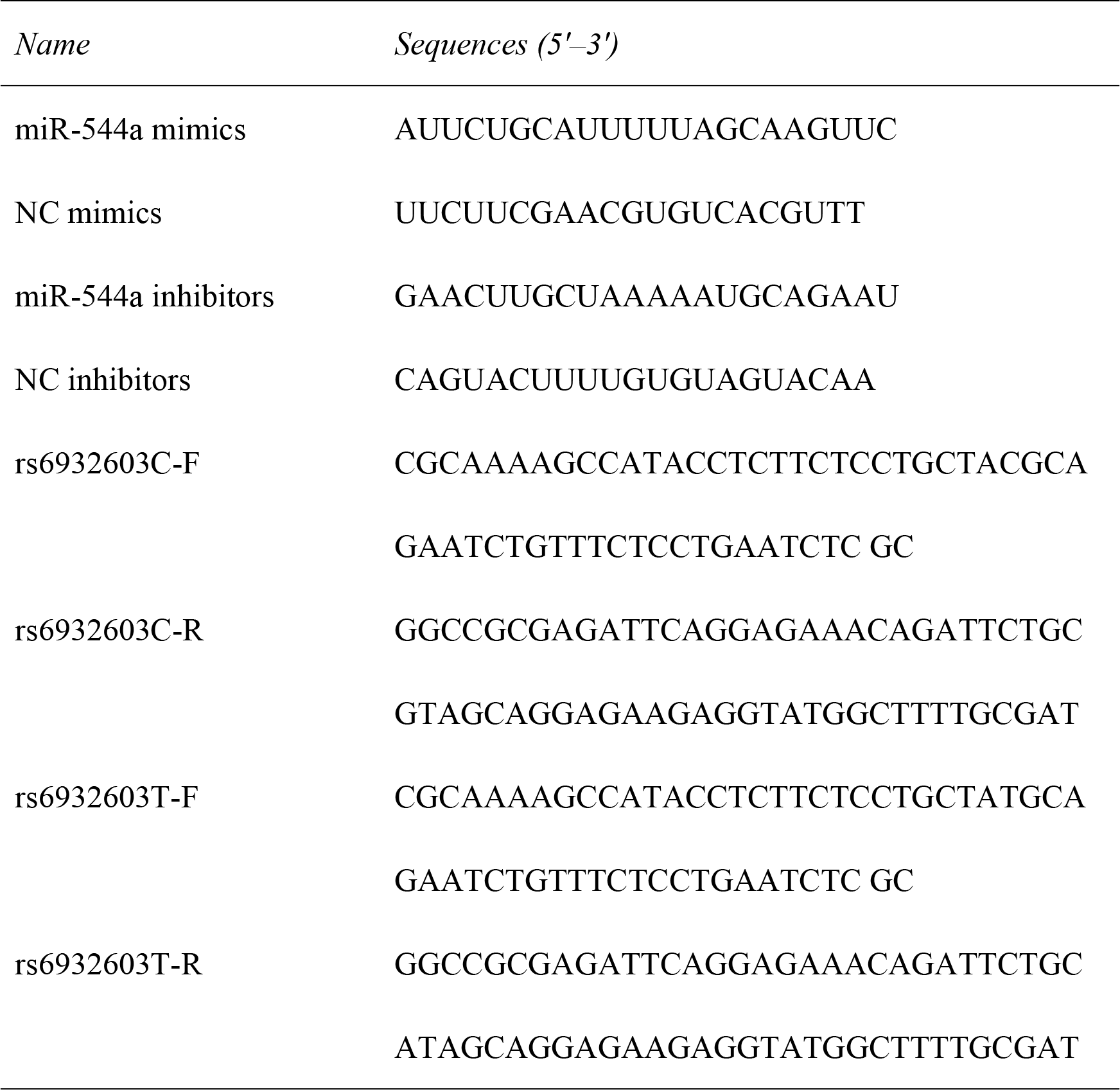
Small fragments of RNA synthesized in the present study

### RNA extraction and real-time polymerase chain reaction (PCR)

Cells were lysed, and total RNA was extracted using TRIzol reagent (Invitrogen) per the manufacturer’s protocol and transcribed into cDNA using ReverAid (Thermo Scientific). Real-time PCR was performed using Hieff qPCR SYBR Green Master Mix (High Rox Plus) (Yeasen). Table 2 lists the PCR primer sequences.

**Table 2.**
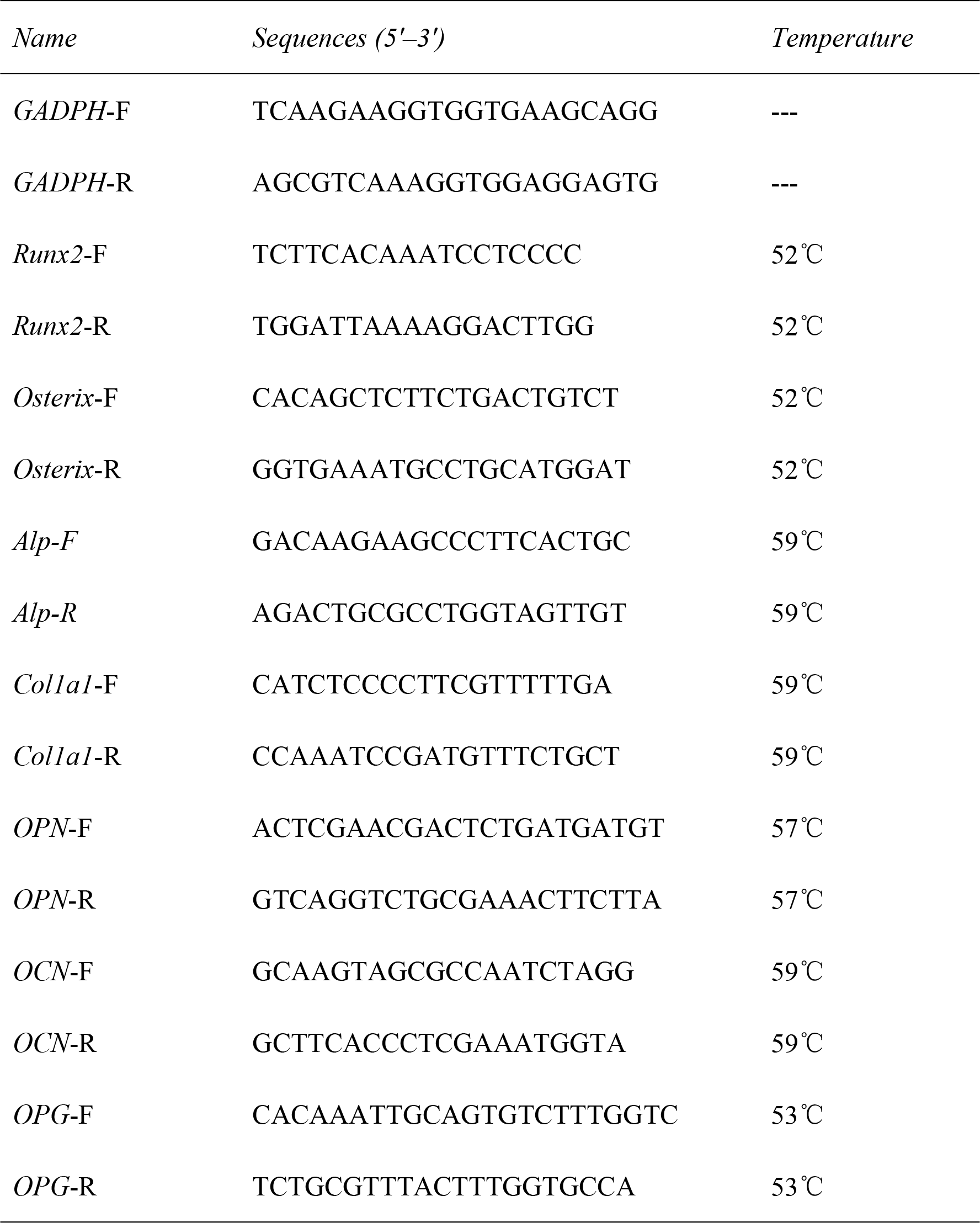
PCR primers sequences in the present study

### Statistical analysis

Data are presented as mean ± SD. Statistical differences between two groups were determined by the twotailed Student’s t test. Statistical differences among groups were analyzed by one-way ANOVA followed by Student–Neuman–Keuls’ multiple comparisons test. All experiments were performed independently at least three times with similar results, and representative experiments are shown. P < 0.05 was considered statistically significant. NS > 0.05, *P < 0.05, **P < 0.01, ***P < 0.001.

## Results

### Differences in rs6932603-C and -T allele expressions

To determine whether the rs6932603-C and rs6932603-T allele expressions differed, 293T and U2OS cells were transfected with the rs6932603-C and rs6932603-T allele psi-check-2 vectors. The luciferase reporter assay showed that the rs6932603-T allele expression level was significantly lower than that of the rs6932603-C allele in the 293T cells and U2OS cells (Figs. 1A and 1B).

**Fig. 1.**
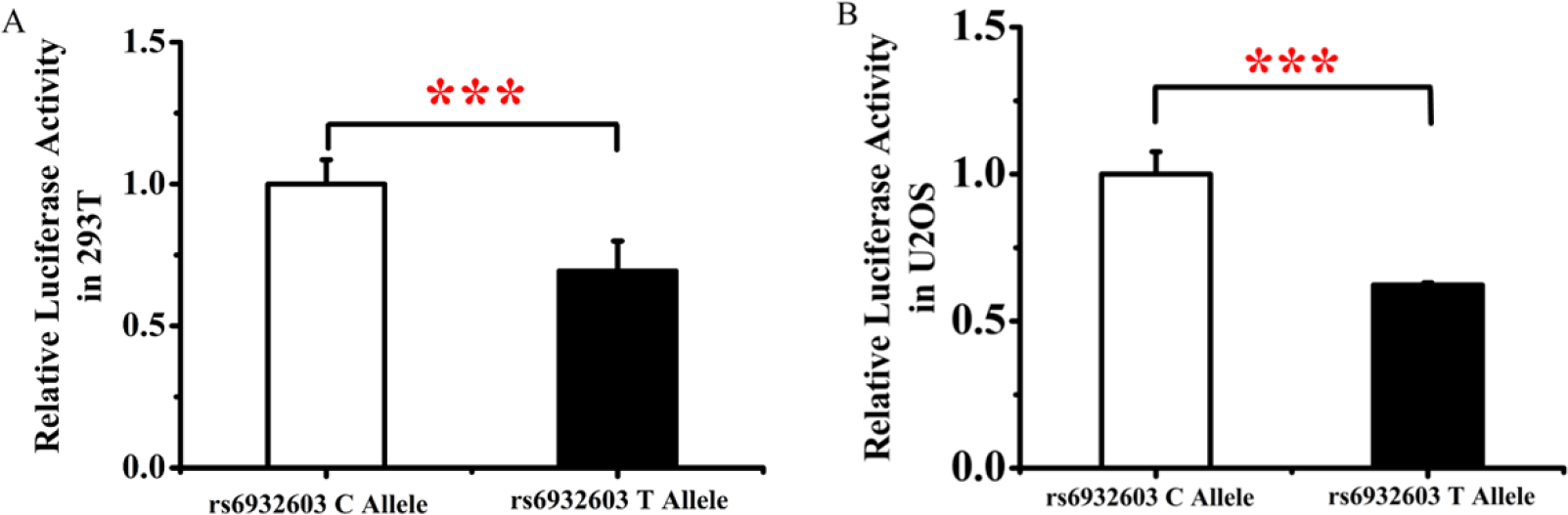
Expression differences between the rs6932603-C and -T alleles. (A) Expression level of the rs6932603-T allele was significantly lower than that of the C allele in 293T cells. (B) Expression level of the rs6932603-T allele was significantly lower than that of the C allele in U2OS cells. Results are shown as percentages relative to luciferase activity. Data are from three independent transfection experiments with assays performed in triplicate (n=6). Luciferase signaling was normalized to Renilla signaling. Error bars show the standard deviation for six technical replicates of a representative experiment. P-values were calculated using a two-tailed Student’s t-test. ***P<0.001.

### MiR-544a differentially regulated the allele variants of rs6932603

Next, we tested a biological model in which miR-544a differentially regulated the C/T allele variants of rs6932603 in *CCDC170*. In this model, rs6932603-T decreased the *CCDC170* transcription levels, leading to an increased osteoporosis risk as predicted (http://mirdsnp.ccr.buffalo.edu/ (Fig. 2A). To test our model, we cotransfected the rs6932603-C/T allele psi-check-2 vector, miR-544a mimics and NC mimics in 293T cells and U2OS cells. The luciferase reporter assay results showed that the miR-544a mimics significantly downregulated rs6932603-T expression in both cell lines (Figs. 2B and 2C). As an additional test of our model, the rs6932603-C/T allele psi-check-2 vector, miR-544a inhibitors and NC inhibitors were cotransfected into 293T cells and U2OS cells. The rs6932603-T expression was significantly upregulated in both cell lines (Figs. 2D and 2E). These results indicated that miR-544a inhibits *CCDC170* expression by differentially binding rs6932603-C/T.

**Fig. 2.**
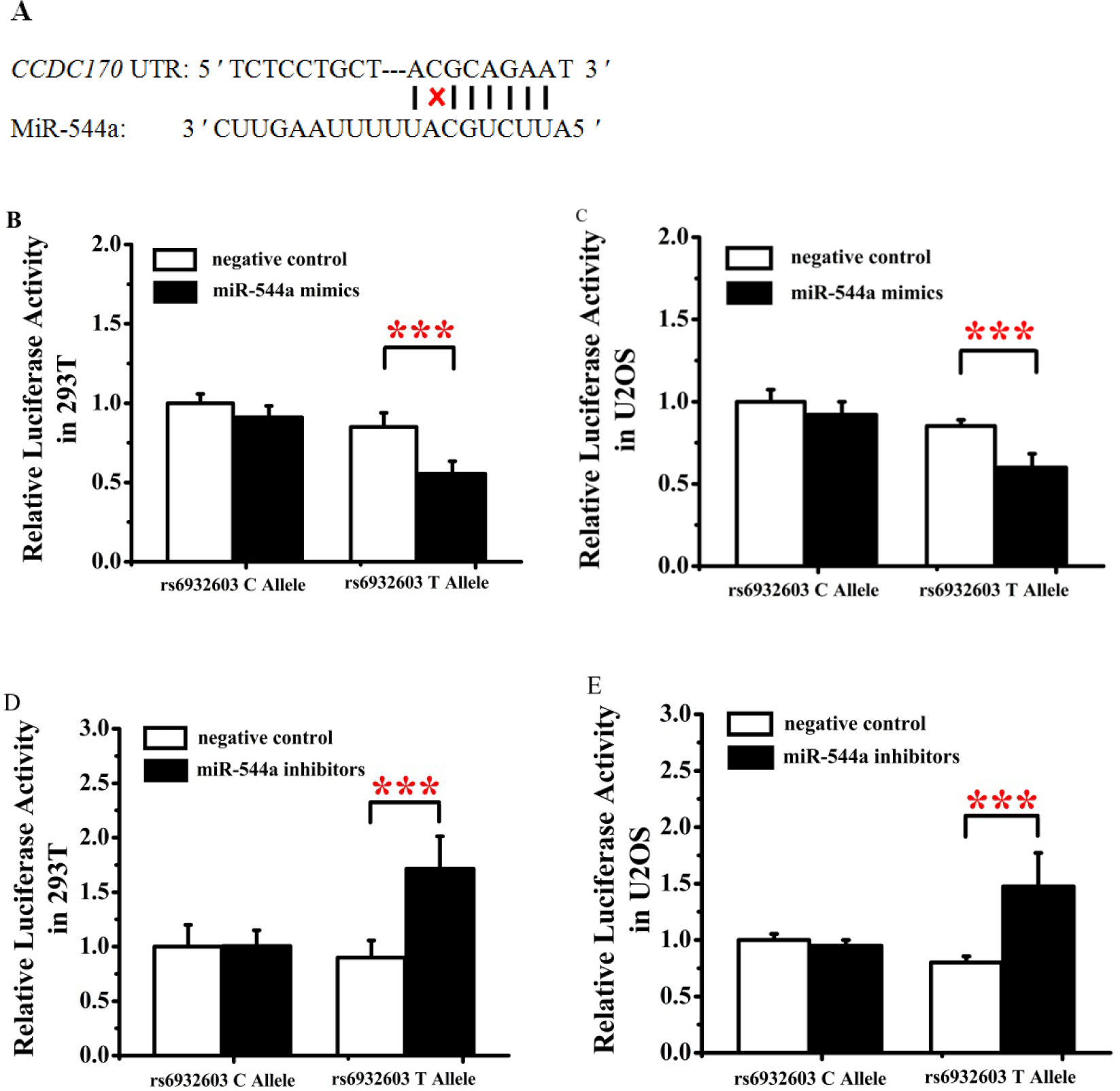
MiR-544a differentially regulates the allele variants of rs6932603. (A) Schematic diagram of the miR-544a binding site on *CCDC170*. (B) Overexpression of miR-544a significantly suppressed rs6932603-T allele levels in 293T cells. (C) Overexpression of miR-544a significantly suppressed rs6932603-T allele levels in U2OS cells. (D) Knockdown of miR-544a significantly upregulated rs6932603-T allele levels in 293T cells. (E) Knockdown of miR-544a significantly upregulated rs6932603-T allele levels in U2OS cells. Results are shown as percentages relative to luciferase activity. Data are from three independent transfection experiments with assays performed in triplicate (n=6). Luciferase signaling was normalized to Renilla signaling. Error bars show the standard deviation for six technical replicates of a representative experiment. P-values were calculated using a two-tailed Student’s t-test. ***P<0.001.

### MiR-544a overexpression repressed the expression of osteogenesis marker genes

To explore the functional role of miR-544a in osteosarcoma and osteoporosis, miR-544a mimics and NC mimics were transfected into U2OS cells. qRT-PCR showed that expressions of the osteogenesis early-stage marker genes, *Runx2* and *Osterix*, the osteogenesis medium-stage marker genes, *Alp* and *Col1a1*, and the osteogenesis late-stage marker genes, *OPN* and *OCN*, were significantly downregulated (Figs. 3A–3F). We also found that osteoclastogenesis inhibitory factor (*OPG*) expression was significantly downregulated (Fig. 3G). These results indicated that miR-544a overexpression inhibited osteogenesis and promoted osteoclastogenesis.

**Fig. 3.**
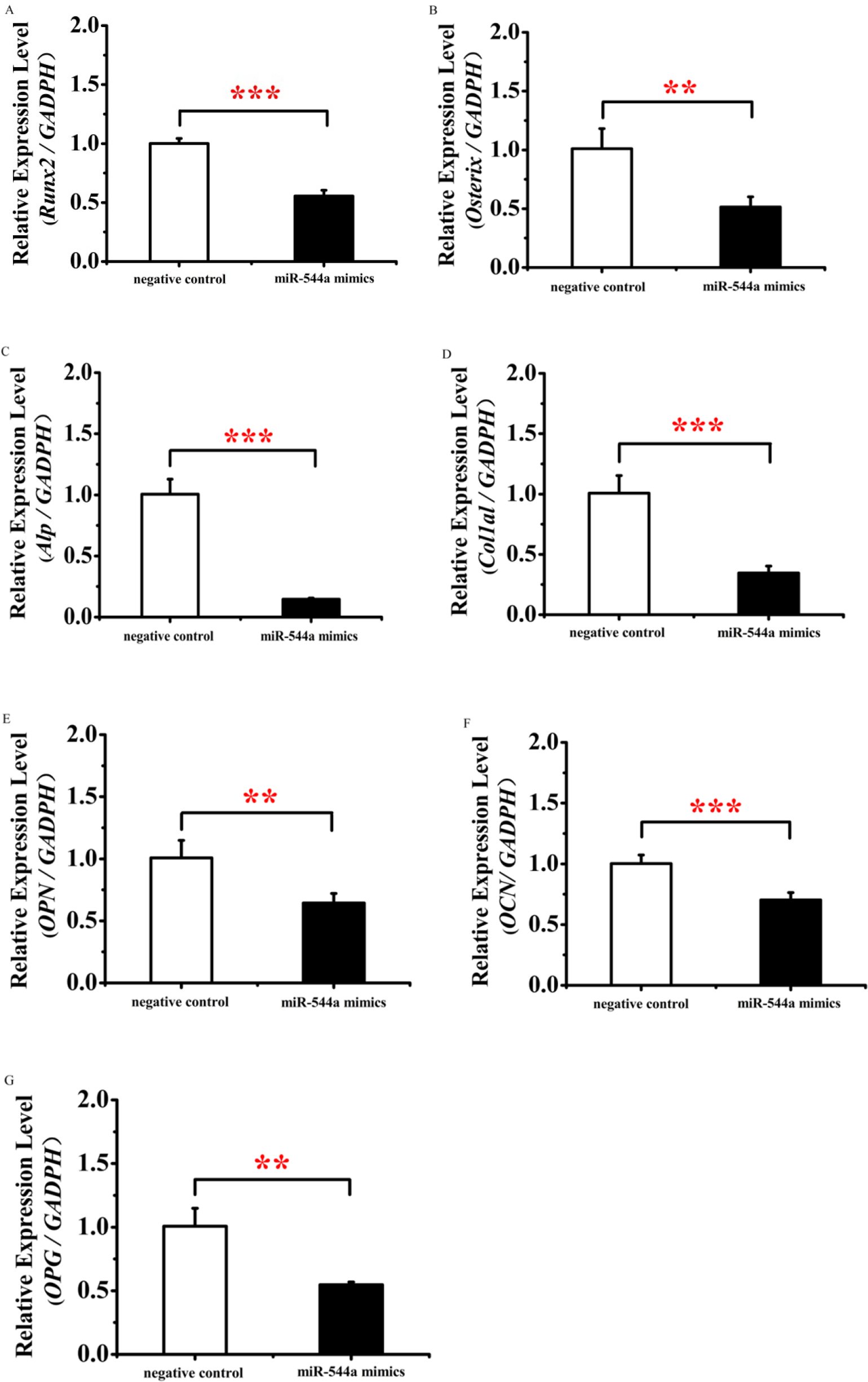
MiR-544a overexpression repressed the expression of osteogenesis marker genes. (A-B) Overexpression of miR-544a significantly suppressed the expression of *Runx2* and *Osterix*. (C-D) Overexpression of miR-544a significantly suppressed the expression of *Alp* and *Col1a1*. (E-F) Overexpression of miR-544a significantly suppressed the expression of *OPN* and *OCN.* (G) Overexpression of miR-544a significantly suppressed the expressions of *OPN* and *OPG*. Data are from three independent transfection experiments with assays performed in triplicate (n=8). Error bars show the standard deviation for four technical replicates of a representative experiment. P values were calculated using a two-tailed Student’s t-test. **P<0.01,***P<0.001.

### MiR-544a knockdown increased the expression of osteogenesis marker genes

To further verify the role of miRNA in inhibiting osteogenesis and promoting osteoclastogenesis, miR-544a inhibitors and NC inhibitors were transfected into U2OS cells. By qRT-PCR, we found that the expressions of *Runx2*, *Osterix*, *Alp*, *Col1a1*, *OPN* and *OCN* were significantly upregulated (Figs. 4A–4F). We also found that *OPG* expression was significantly upregulated (Fig. 3G). These results indicated that knockdown-expression of miR-544a promoted osteogenesis and inhibited osteoclastogenesis.

**Fig. 4.**
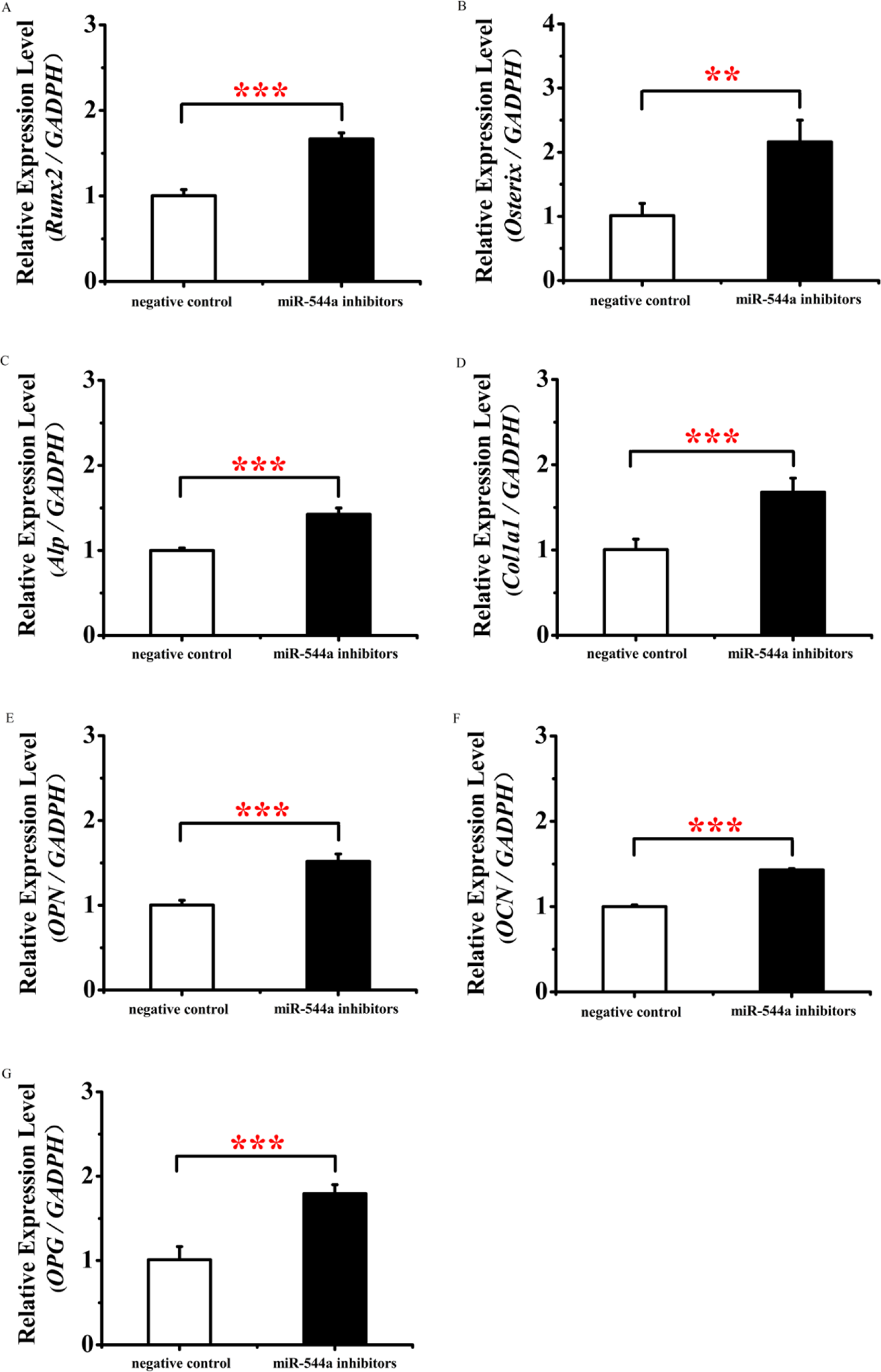
MiR-544a knockdown increased the expressions of osteogenesis marker genes. (A-B) Knockdown-expression of miR-544a significantly increased the expressions of *Runx2* and *Osterix*. (C-D) Knockdown-expression of miR-544a significantly increased *Alp* and *Col1a1* expression. (E-F) Knockdown-expression of miR-544a significantly increased the expressions of OPN and OCN. (G) Knockdown-expression of miR-544a significantly increased *OPG* expression. Data are from three independent transfection experiments with assays performed in triplicate (n=8). Error bars show the standard deviation for four technical replicates of a representative experiment. P-values were calculated using a two-tailed Student’s

## Discussion

Identifying target genes is pivotal to understanding miRNA’s roles in various diseases, including osteoporosis[17]. The TargetScan, miRNASNP and miRbase databases were used to identify direct targets of miR-544a. *CCDC170* was predicted as one target of miR-544a. In our study, we found that miR-544a differentially regulated the allele variants of rs6932603-C/T in the *CCDC170* 3′-UTR via the luciferase reporter assay. Our study provided a possible mechanism for this, in that upregulation of miR-544a resulted in decreased *CCDC170* expression in osteoporosis. Multiple studies have found that *CCDC170* plays an important role in breast cancer [18,19] and multiple genome-wide linkage scans have shown that *CCDC170* is one of the genes most strongly linked to BMD. However, no in vivo or in vitro experiments have directly verified the role of *CCDC170*. We plan to study this in our next research project.

MicroRNAs influence posttranscriptional regulation in various genes, and some have been shown to influence osteoporosis[20,21]. MiR-544a has been shown to play an important role in many diseases such as lung cancer and breast cancer, but its role in osteoporosis is unknown[22,23]. Here, we showed that miR-544a suppressed osteogenesis and promoted osteoclastogenesis by downregulating the expressions of *RUNX2*, *Osterix*, *ALP, Col1a1*, *OPN*, *OCN* and *OPG*.

In conclusion, our study demonstrated that miR-544a inhibits *CCDC170* expression by differentially binding rs6932603-C/T. MiR-544a also suppresses osteogenesis and promotes osteoclastogenesis by downregulating the expressions of osteogenesis marker genes. These findings may improve our understanding of the association between miRNAs, GWAD lead SNPs, and osteoporosis pathogenesis and may provide a potential therapeutic target for osteoporosis therapy.

